# Spatially aware self-representation learning for tissue structure characterization and spatial functional genes identification

**DOI:** 10.1101/2023.03.13.532390

**Authors:** Chuanchao Zhang, Xinxing Li, Wendong Huang, Lequn Wang, Qianqian Shi

## Abstract

Spatially resolved transcriptomics (SRT) enable the comprehensive characterization of transcriptomic profiles in the context of tissue microenvironments. Unveiling spatial transcriptional heterogeneity needs to effectively incorporate spatial information accounting for the substantial spatial correlation of expression measurements. Here, we develop a computational method, SpaSRL (spatially aware self-representation learning), which flexibly enhances and decodes spatial transcriptional signals to simultaneously achieve spatial domain detection and spatial functional genes identification. This novel tunable spatially aware strategy of SpaSRL not only balances spatial and transcriptional coherence for the two tasks, but also can transfer spatial correlation constraint between them based on a unified model. Additionally, this joint analysis by SpaSRL deciphers accurate and fine-grained tissue structures and ensures the effective extraction of biologically informative genes underlying spatial architecture. We verified the superiority of SpaSRL on spatial domain detection, spatial functional genes identification and data denoising using multiple SRT datasets obtained by different platforms and tissue sections. Our results illustrate SpaSRL’s utility in flexible integration of spatial information and novel discovery of biological insights from spatial transcriptomic datasets.

## Introduction

Recent advances in spatially resolved transcriptomics (SRT) have enabled high-throughput sequencing of mRNA coupled with spatial information in multicellular organisms, which can resolve cellular localizations to unveil the organizational landscape of complex tissues [1]. The SRT sequencing-based techniques, such as Spatial Transcriptomics (ST) [2], 10x Visium, Slide-seqV2 [3], can measure the expression level of tens of thousands of genes in thousands of tissue locations (or spots), which enables the comprehensive study of spatial transcriptional landscape from tissue architecture heterogeneity and the corresponding functional genes. However, SRT measurements are often sparse and noisy due to various technical limitations, e.g., transcript capture rate or spatial resolution, which pose great challenges to decipher the spatially functional regions and genes [4]. Apart from dimension reduction, an effective usage of the locational information contained in SRT data can also mitigate data noise or bias, improving the pattern recognition in SRT studies, since neighboring locations on tissue often share cell microenvironments and display similar gene expression levels in spatial transcriptomics [5].

To resolve tissue structure, spatial domain detection is an important research topic, which aims to cluster spots with similar gene expression and spatial continuity within each cluster (or spatial domain). For this purpose, several currently presented spatial clustering approaches, e.g., BayesSpace [6], Hidden Markov Random Field (HMRF) [7], SEDR [8], STAGATE [9] and SpaGCN [10], additionally constrain the models with spatial information to facilitate the identification of spatial domains with spatial smoothness. Their outcomes display more spatially continuity than those from clustering methods previously developed for single-cell RNA-sequencing (scRNA-seq) studies that only utilize expression measurements, e.g., Seurat [11], SCANPY [12]. However, these methods often take spatial neighbor prior as a hard constraint to ensure spatial continuity in spatial domains, but seldom provide a flexible solution to balance spatial coherence and expression variability within neighborhoods.

In addition, most existing methods substantially perform dimension reduction before clustering spots and the common approach is principal component analysis (PCA) that is adopted to preprocess SRT data by e.g., Seurat, SCANPY, BayesSpace, SEDR and SpaGCN. PCA or other linear dimension reduction techniques can not only mitigate data noise but also extract the potential functional genes from co-expression or functional association perspective [13]. But this kind of usage of dimension reduction for SRT studies does not ensure the inferred components (or meta genes) relevant to the spatial map on tissue, thus may reducing their effectiveness or biological interpretations. Whereas, some recent works have made efforts to deal with this issue. For example, SpatialPCA [5] incorporates spatial information to improve locational neighborhood similarity in the constructed PC space. DR-SC [14] performs simultaneous clustering and dimension reduction for better biological associations between the detected clusters and (meta) genes. However, it is still challenging to unify the characterization of locational and gene patterns accounting for spatial coherence and biological interpretations in SRT studies.

To this end, we present **sp**atially **a**ware **s**elf-**r**epresentation **l**earning (SpaSRL), a novel method that achieves spatial domain detection and dimension reduction in a unified framework while flexibly incorporating spatial information. Specifically, SpaSRL enhances and decodes the shared expression between spots for simultaneously optimizing the low-dimensional spatial components (i.e., spatial meta genes) and spot-spot relations through a joint learning model that can transfer spatial information constraint from each other. SpaSRL can improve the performance of each task and fill the gap between the identification of spatial domains and functional (meta) genes accounting for biological and spatial coherence on tissue. Thus, SpaSRL not only deciphers fine-grained spatial domains and extracts spatial interpretable functional genes underlying spatial domains, but also corrects the low-quality gene expression from borrowing information within spatial clusters, which flexibly balances spatial coherence and expression variability.

We demonstrate the superiority of SpaSRL to identify accurate spatial domains and functional (meta) genes on datasets sequenced by different technologies. We illustrate that SpaSRL can flexibly exert spatial information on the identification of spatial domains and functional genes as complementary to the current usage of spatial information. Applied to breast cancer slices, SpaSRL deciphers intratumor heterogeneity and finds more novel cancer associated genes, which are validated by the survival analysis of independent clinical data. Applied to brain slices, SpaSRL reveals the tissue structures and the corresponding functional genes for interpreting tissue functions.

## Methods

### Overview of SpaSRL

SpaSRL first enhances the shared expression among spots by incorporating spatial information into gene expression (Figure 1A). SpaSRL borrows the shared information from spatially neighboring spots to adjust gene expression in each spot (i.e., original expression matrix *X*^0^ ∈ *R*^*M*×*N*^ → enhanced expression data *X* ∈ *R*^*M*×*N*^, *M* and *N* respectively denote the number of genes and spots), which can correct the low-quality expression measurements and enrich local signals. The parameter *α* is set to flexibly control the spatial information constraint on gene expression measurements.

**Figure 1.**
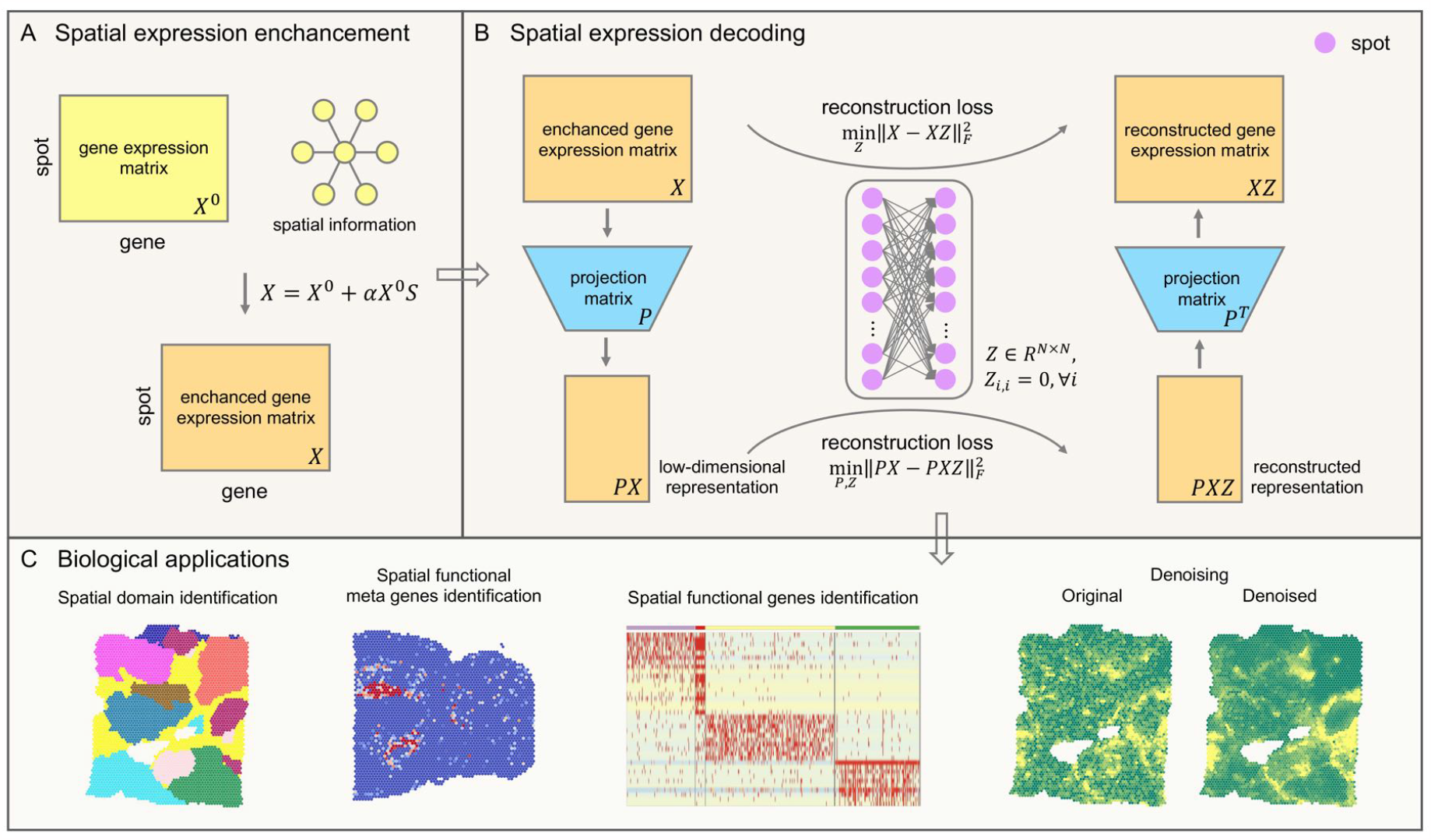
Schematic overview of SpaSRL enhancing-decoding (A&B) processes and potential applications of SpaSRL in downstream SRT analysis (C). (A) Spatial expression enhancement from aggregating expression information from neighborhood spots. SpaSRL incorporates spatial information into gene expression to enhance the shared expression between spots by flexibly aggregating the weighed gene expression from their *k* spatial neighbors (e.g., *X*^0^ → *X*). *S* denotes the weight of expression similarity between each spot and its *k* neighbors. *α* controls the contribution of spatial similarity to the enhanced expression measurements. (B) Spatial expression decoding via the feature extraction embedded self-representation learning model. The input data (i.e., *X*) is the enhanced gene expression matrix from (A). SpaSRL uses a robust projection matrix (i.e., *P*) to generate the low-dimensional representation (i.e., *X* → *PX, P*^*T*^*PX* = *X*). Based on the original and low-dimensional data, SpaSRL performs data reconstructions via an aggregated weight matrix *Z* based on self-representation learning algorithm (i.e., *X* = *XZ* and *PX* = *PXZ*). SpaSRL iteratively learns the projection matrix (i.e., *P*) and spot-spot similarity matrix (i.e., *Z*) by minimizing the sum of reconstruction losses (see Methods). When SpaSRL reaches convergence, the two optimal matrices are achieved for further downstream analyses. (C) Biological applications for SpaSRL including spatial domain identification, functional genes/meta genes identification and data denoising. The spot-spot similarity matrix can be applied to detect spatial domains and data denoising. The projection matrix can be employed to identify functional genes/meta genes to improve biological insights into tissue heterogeneity.

SpaSRL then decodes the spatially enhanced biological signals based on a novel self-representation learning model (Figure 1B). In the model, SpaSRL introduces an aggregated weight matrix *Z* ∈ *R*^*N*×*N*^ and a projection matrix *P* (i.e., *P* ∈ *R*^d×*M*^, *d* ≪ *M, d* is the number of meta genes) to reconstruct expression data as the linear combinations of expression measurements of similar locations in the original and low-dimensional feature spaces (i.e., *X* → *PX, X* = *XZ* and *PX* = *PXZ*). The aggregated weight matrix *Z* measures the contribution of other spots to each spot, which reflects the transcriptional and neighborhood similarity structure of spots. To achieve the robust projection matrix *P*, the matrix *P*^*T*^ should satisfy the constraint of restoring the original expression data (i.e., *P*^*T*^*PX* = *X*). By minimizing the reconstruction errors, SpaSRL iteratively updates *Z* and *P* by fixing the other to obtain the optimal solutions.

The optimal *Z* and *P* can be used for multiple downstream analytical tasks (Figure 1C). The optimal *Z*, denoted as the spot-spot similarity matrix, enables SpaSRL to (1) detect spatial domains interoperating with Leiden [15] or Louvain [16] methods, and serves to (2) denoise expression profiles (i.e., *XZ*) to improve gene spatial expression patterns for individual gene analysis, e.g., spatially variable or differentially expressed genes identification. The optimal *P*, as the spatial component loading matrix, stores potential meta genes fitting in with the spatial neighboring structure, whereby enabling SpaSRL to (3) extract the spatial functional gene sets relevant to different spatial domains (see Methods).

The primary advantage of SpaSRL is the joint solution of *Z* and *P* with spatial information constraint, which can transfer the spatially enriched biological signals between samples (spots) and genes, resulting in the robust identification of spot clusters and spatial functional (meta) genes with refined spatial patterns and favorable biological associations. Additionally, SpaSRL provides a unique spatially aware strategy that allows tunable usage of spatial information, which not only can, to a great extent, correct for dropouts or noises, but also can control the impact of spatial neighborhood similarity on spatial domains and functional meta genes by tuning parameter *α* manually or according to the obtained outcomes. SpaSRL method, altogether with the landmark-based strategy provided for large-scale datasets, is computationally optimized. Furthermore, we distribute SpaSRL as a user-friendly Python module based on the widely used AnnData data structure.

### Constructing spatially aware self-representation learning model

#### Enhancing the shared expression between spots

We incorporate spatial information into gene expression to enhance the shared expression between spots, which can correct the low-quality gene expression (e.g., dropout) in each spot by borrowing information from its surrounding neighborhood. Using a weight matrix *S* ∈ *R*^*N*×*N*^, the original expression matrix *X*^0^ is adjusted as the enhanced expression data *X* specifically as follows:

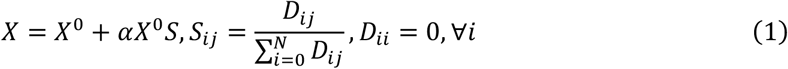

where the similarity matrix *D* is obtained by calculating the cosine distances between *k*-nearest spatial neighbor spots on top 15 principal components (i.e., *D* = exp(2 − *cosine*_*dist*(*U*)), *U* ∈ *R*^15×*N*^). The default number of neighbors, i.e., *k*, is set to 10 for 10x Visium datasets and 30 for Silde-seqV2 datasets in this work. The tunable parameter *α* can be flexibly set, which controls the extent to aggregating expression across surrounding spots for generating the enhanced expression data *X*. When the value is set to 1, the spot itself and spatial neighbors contribute equally to the enhanced data.

#### Decoding the shared expression between spots

We build a feature extraction embedded self-representation learning model to reconstruct data in original and low-dimensional latent spaces, which decodes the enhanced expression to measure the contribution of other spots to each spot, enabling the optimalization of low-dimensional spatial components and spot-spot relations. The main procedure can be stated as follows.

- *Data reconstruction of the enhanced data*: Suppose there is an aggregated weight matrix *Z* ∈ *R*^*N*×*N*^ (i.e., spot-spot similarity), SpaSRL reconstructs the enhanced gene expression *X* ∈ *R*^*M*×*N*^ by aggregating the shared gene expression across spots with the Frobenius norm:

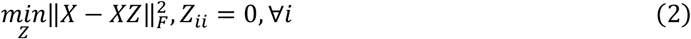
- *Data reconstruction of the low-dimensional representation*: To mitigate data dropouts and extract robust spatial components (or meta genes), SpaSRL leverages a projection matrix *P* to generate the low-dimensional representation (i.e., *PX*) and further to restore the enhanced data (i.e., *P*^*T*^*PX* = *X*). In consideration of the consistency of spot-spot similarity in both original and low-dimensional spaces, the aggregated weight matrix *Z* also needs to satisfy the reconstruction of the low-dimensional representation. Note that, *l*_2,1_-norm should be used to replace *F*-norm and further to improve the ability to simultaneously learn dimension reduction and the relationship between samples due to the existence of constraint terms (i.e., *P*^*T*^*PX* = *X*).

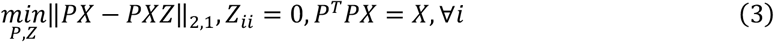

Clearly, the main objective function can be bluntly written as the combination of the above terms in Eq.(4), by which we can solve the optimal *Z* as the spot-spot similarity and the optimal *P* as the spatial components.

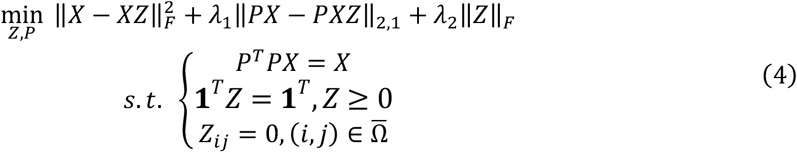

where **1** represents an all-one vector for normalization. 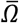 is the complement of *Ω* which is a set of connections of samples (spots) in an adjacency graph. If 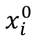 and 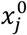 are not connected in the adjacency graph, then we have 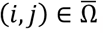. The adjacency graph is determined by *K*-nearest neighbor (*K*NN) algorithm with Euclidean distances of all samples. Parameter *K* may be chosen freely and there are two tunable parameters *λ*_1_ and *λ*_2_ to balance the three terms in Eq. (4). Both parameters can be determined according to data properties or settled empirically. We discussed the sensitivity of SpaSRL to these parameters in Supplementary Figure 1 and proved that the clustering performance of SpaSRL is robust in a large range of *K, λ*_1_ and *λ*_2_.

Note that, SpaSRL is a variant of Low-Rank Representation learning (or self-representation learning), whose standard penalty term should be the nuclear-norm (i.e., ‖*Z*‖_∗_). The nuclear-norm can constrain the matrix *Z* to have a better cluster structure [17], but relies on eigenvalue decomposition operator in the solving process, thus will greatly increasing the running time of SpaSRL. To optimize the computational efficiency, we use the F-norm instead due to the existing relations between the two norms (i.e., ‖*Z*‖_*F*_ ≤ ‖*Z*‖_∗_). Additionally, to discuss the necessities of each component in loss function, the ablation experiment is performed on the benchmark datasets (Supplementary Figure 2). The ablation experiment indicates that each component of the loss function is necessary and further confirms the effectiveness of the loss function design.

#### Solving spatially aware self-representation learning model

The spatially aware self-representation learning model (i.e., Eq. (4)) presents as a linear-equality constrained problem, which can be solved by the alternating direction method of multipliers (ADMM) [18]. Thus, Eq. (4) can be equivalently transformed to:

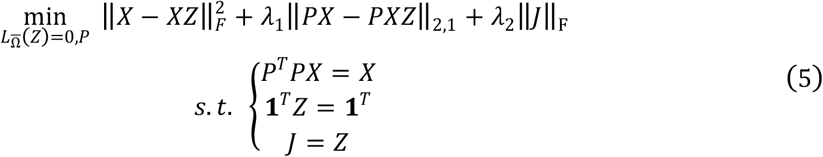

where 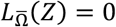 corresponds to the third constraint in Eq. (4). Then, the augmented Lagrangian function of Eq. (5) is:

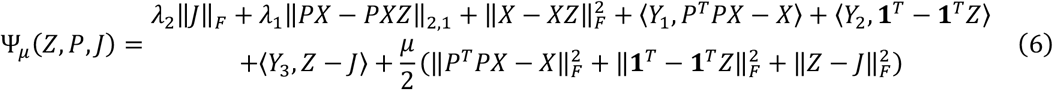

where *μ* denotes a penalty parameter larger than 0. ‖·‖_*F*_ represents the Frobenius norm. *Y*_1_, *Y*_2_ and *Y*_3_ are the corresponding Lagrangian multipliers in Eq. (6). Thus, the above problem becomes unconstrained, and according to ADMM algorithm, it can be minimized in turn to update the variables *Z, J, P* with the other variables fixed.

Specifically, supposing that after *k* times of updates with *Z*^*k*^, *J*^*k*^ and *P*^*k*^, the next update at iteration *k* + 1 can be written as:

1. Solving the optimal matrix *J* of Eq. (5) with all other matrices fixed

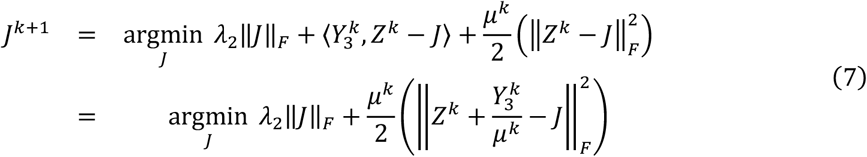
2. Solving the optimal matrix *Z* of Eq. (5) with all other matrices fixed

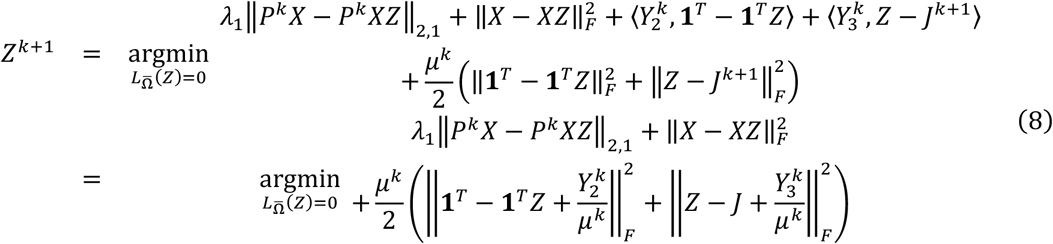
3. Solving the optimal matrix *P* of Eq. (5) with all other matrices fixed

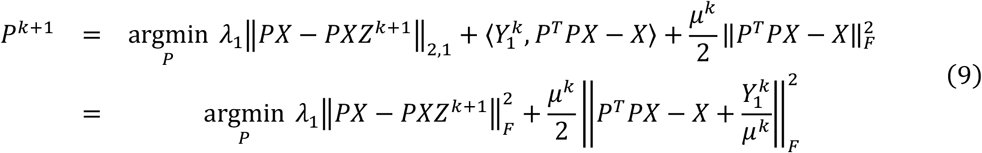

The optimal solutions of *Z* and *P* are obtained by iteratively solving the subproblems 1)-3) until convergence. For better clarity, the corresponding pseudocode of main solving process is summarized in Algorithm 1 in Supplementary Note S1.

#### Landmark-based spatially aware self-representation learning for large-scale datasets

When dealing with large-scale datasets (e.g., tens of thousands of samples [spots] or more), SpaSRL will consume a lot of times and storage to build the similarity matrix between all samples or spots (i.e., *Z* ∈ *R*^*N*×*N*^). To improve the capacity of SpaSRL on large-scale datasets, we propose a landmark-based strategy to facilitate the widespread application of SpaSRL on different SRT platforms. The key idea of landmark-based strategy is to select a small number of samples that should be representatives of the underlying sample manifold and then construct a landmark-by-sample matrix (i.e., *V* ∈ *R*^*L*×*N*^, *L* ≪ *N, L* is the size of landmark sample set) to approximate the original sample-by-sample matrix (i.e., *Z* ∈ *R*^*N*×*N*^). Since the number of landmarks is much smaller than the total number of samples, this approximation can significantly reduce memory and time occupation. SpaSRL uses *K*-means method to select these landmarks, which are the samples nearest to the real cluster center. This landmark-based approximation strategy is inspired by a previous work [19], which can theoretically ensure the effectiveness for large-scale datasets.

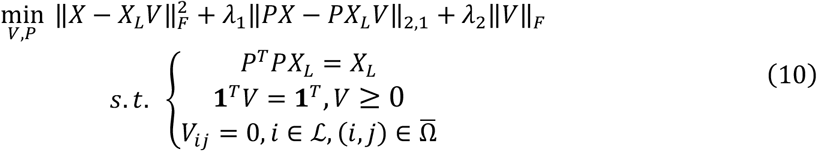

where *X*_*L*_ ∈ *R*^*M*×*L*^ is the expression matrix of landmark samples. ℒ is the landmark samples set.

Pseudocode of the landmark-based spatially aware self-representation learning is summarized in Algorithm 2 of Supplementary Note S1. This strategy is recommended to deal with datasets with more than 10,000 samples (e.g., Slide-seqV2 datasets). We discussed the sensitivity of SpaSRL to the number of selected landmark samples using a Slide-seqV2 dataset (contain 39,496 spots) and show that the domains identified by SpaSRL is robust in Supplementary Figure 1.

### Data collection and general preprocessing

There are 17 datasets from 2 different SRT platforms of diverse resolutions in this paper including 14 10x Visium brain datasets, 2 10x Visium breast cancer datasets, and 1 Slide-seqV2 cerebellum data. We firstly selected highly variable genes (HVGs) by using scanpy.pp.highly_variable_genes() from SCANPY Python package [12]. We used the top 3,000 HVGs for 10x Visium datasets and Slide-seqV2 datasets. Then, we performed log-transformation on the expression profiles via scanpy.pp.log1p(), and the transformed data subsequently served as the input of SpaSRL.

### Spatial domain identification and visualization

SpaSRL uses the spot-spot similarity matrix *Z* to identify spatial domains by Louvain [16] or Leiden [15] algorithms, which are respectively implemented as scanpy.tl.louvain() or scanpy.tl.leiden(). For large-scale dataset, the landmark-sample similarity matrix *V* is learned and SpaSRL first uses scanpy.pp.pca(), scanpy.pp.neighbors() and then performs spatial domains detection by Louvain or Leiden. The parameter ‘resolution’ in the functions is adjusted to match the number of annotated structures provided by the original authors or manually defined with prior (anatomical) knowledge. In our practice, we use Louvain for 10x Visium datasets and Leiden for large-scale datasets (i.e., with > 10,000 spots) to identify spatial clusters.

SpaSRL adopts UMAP (Uniform Manifold Approximation and Projection) for spot embedding visualization based on the spot-spot similarity matrix *Z*. When based on the landmark-sample similarity matrix *V* for large-scale dataset, the algorithm first uses scanpy.pp.pca(), scanpy.pp.neighbors() and then performs scanpy.tl.umap() for visualization.

### Gene expression denoising

SpaSRL uses the captured spot-spot similarity matrix *Z* to denoise the gene expression profiles (i.e., *XZ*). For large-scale dataset, SpaSRL uses the captured landmark-sample similarity matrix *Z* and the expression matrix *X*_*L*_ of landmark samples to perform data denoising (i.e., *X*_*L*_*V*).

### Spatial functional genes identification

SpaSRL ranks the weight of each gene on each spatial component in descending order by using matrix *P*. Then, top 500 weighted genes on the spatial components are selected, and the intersection was taken with the differentially expressed genes (i.e., log fold change [LFC ≥ 1]) in each spatial domain from denoised data as specific functional genes of each spatial domain. The differentially expressed genes of each spatial domain are identified via FindAllMarkers() in Seurat R package.

The genes with the top weight from spatial components always show good co-expression properties or functional associations, which can provide better biological interpretations than individual differentially expressed genes. Therefore, we regard the intersection between the top weighted genes and the differentially expressed genes as spatial functional genes, which can elucidate more biologically meaningful and domain-specific features underlying tissue structure.

### Performance evaluation

We describe below the metrics used in this work to evaluate the performance of SpaSRL in two aspects: (1) spatial domain detection, (2) spatial functional genes identification. Details of the benchmarking approaches are provided in Supplementary Note S1.

#### Accuracy and spatial coherence of spatial domains

1) If ground truth annotations are available (e.g., from original publications), adjusted Rand index (ARI) [20] and cluster purity (i.e., Eq. (11)) [6] are used to quantify the accuracy of spatial domain. 2) Local inverse Simpson’s Index (LISI) [21] and Moran’s I statistics [22] are used to quantify the spatial coherence of domains. The LISI value for every sample is computed by using compute_lisi() in lisi R package. The function parameter ‘perplexity’ is set to 10 for 10x Visium datasets and 30 for Slide-seqV2 datasets. The Moran’s I statistics for every spatial domain are computed to measure the spatial autocorrelation via moranI() in Rfast2 R package. For computing the Moran’s I value of each spatial domain, we set the feature vector of the samples belonging to this domain to 1 and other samples to 0, and the weight uses the inverse of Euclidean distance on 2D spatial coordinates of spots.

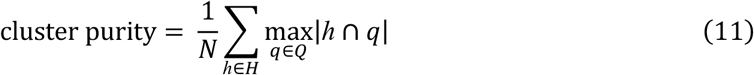

where *H* is the set of clusters set or spatial domains and *Q* is the set of reference groups. Cluster purity is an external evaluation measures of clustering results and measures the extent to which a cluster contains the entities from only one partition. Cluster purity is specifically used to evaluate the clustering performance on spatially resolved transcriptomics (SRT) datasets with rough annotations (e.g., BC and IDC breast cancer slices) [6].

#### Spatial continuity and expression specificity of functional genes

The Moran’s I is used to evaluate the spatial autocorrelation of gene expression before and after denoising. We evaluate gene expression specificity before and after denoising by comparing the LFC values of top marker genes for each domain.

### Survival analysis

We evaluate the prognostic significance of a gene using bulk expression profiling data with patient survival information in breast cancer study. We obtain Breast Cancer International Consortium (METABRIC) breast cancer cohort 1 dataset (*n* = 997 patients) in RTNsurvival R package [23]. Then, we stratify the subjects into high and low groups by using the feature median value (i.e., gene expression), and perform Kaplan Meier (KM) analysis between the two groups to compare the survival difference.

### Functional/cancer hallmark enrichment analysis

The R package clusterProfiler [24] is used for functional/cancer hallmark enrichment analysis of the discovered functional genes.

## Results

### Benchmark the performance of SpaSRL on revealing tissue structures and tumor heterogeneity

We quantitively evaluated the ability of SpaSRL to identify spatial domains using 14 10x Visium arrays, including 12 human dorsolateral prefrontal cortex (DLPFC) slices and 2 breast cancer slices. We took the manual annotation of DLPFC slices provided by the original authors [25] as ground truth and quantified the similarity between identified clusters and the manual labels using adjusted Rand index (ARI). We benchmarked SpaSRL against existing spatial (i.e., BayesSpace [6], Giotto [7], SEDR [8], SpaGCN [10], stLearn [26], STAGATE [9] and Vesalius [27]) and non-spatial (i.e., non-negative matrix factorization [NMF] [28], self-representation [17], variational autoencoder [VAE] [29], Leiden implemented in SCANPY [12] and Louvain implemented in Seurat [11]) clustering methods. Overall, SpaSRL achieved the highest mean ARI (mean ARI = 0.54) and substantially outperformed the competing methods (Wilcox signed rank test, *P* < 10^−6^, Figure 2A). Moreover, SpaSRL had obvious advantage in time efficiency over most of the involved methods (Wilcox signed rank test, *P* < 10^−5^, Figure 2B). The results also show that spatial clustering methods generally perform better than non-spatial methods (Wilcoxon signed-rank test, *P* < 10^−4^, Figure 2A), indicating that the usage of spatial information can improve the identification of spatial domains.

**Figure 2.**
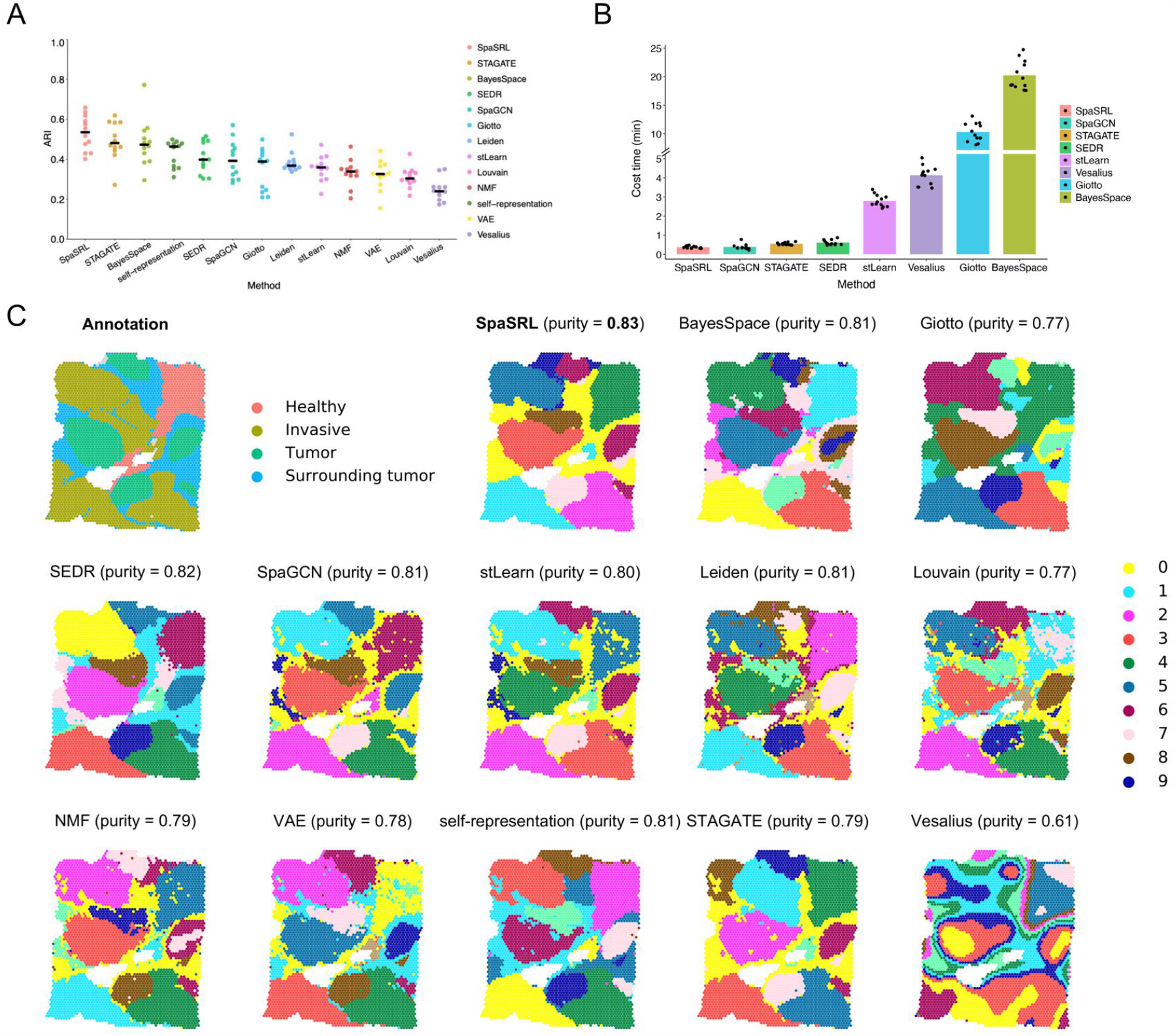
Comparative performance of SpaSRL to existing spatial and non-spatial methods on spatial domain identification. Summary of clustering performance on 12 manually annotated spatialLIBD datasets in terms of ARI values (A) and time consumption (B). Each point denotes the measured performance on one dataset. The center line in (A) indicates the mean ARI value of each method on all datasets. The methods in (A) are ordered by decreasing mean ARI values. The height of bar in (B) indicates the mean running time of each method on all datasets. The methods in (B) are ordered by increasing mean running time. (C) The comparison of spatial domain identification on BC (*n* = 3,798 spots) slice. The histopathological annotation is obtained from original work [8] and used to color each spot in spatial coordinates without the H&E-stained image. Spatial domains identified by SpaSRL and competing methods on BC slice are distinguished by colors without strict correspondence. The cluster purity is used to compare the similarity between identified outcomes and the reference annotation.

In addition, we assessed these methods for detecting tumor heterogeneity across different breast cancer slices (i.e., 10x Visium Human Breast Cancer Block A Section 1 [BC] and Invasive Ductal Carcinoma [IDC]). We took the histopathological annotations [6, 8] as reference while more clusters reflecting potential transcriptional heterogeneity can be revealed by computational methods (Figure 2C and Supplementary Figure 3). We also benchmarked these spatial and non-spatial clustering approaches by computing cluster purity. Among these methods, SpaSRL achieved the highest cluster purity (purity = 0.82 in BC and purity = 0.87 in IDC) and detected more biologically homogenous structure than other involved methods (Figures 3B, 4A, C and Supplementary Figure 4). Moreover, the spatial domains identified by SpaSRL exhibited better spatial coherence than the outcomes from stLearn, SpaGCN and all the non-spatial clustering methods (i.e., Leiden, Louvain, NMF, VAE and self-representation) as they identified many scattered noisy subclusters (Supplementary Figures 5-6). Other spatial clustering methods also have good spatial continuity in spatial domains, but some (e.g., Giotto and Vesalius) undetected the refined boundaries of certain domains, thus reducing their clustering performance. Additionally, we verified the universality of SpaSRL in quantitative or qualitative ways based on whether data annotation is provided (e.g., Slide-seqV2 [30], Seq-Scope [31], 4i and MIBI-TOF [32]) (Supplementary Figures 7-9). These benchmark tests demonstrated the superiority of SpaSRL at identifying spatial functional domains accounting for spatial coherence and biological difference.

**Figure 3.**
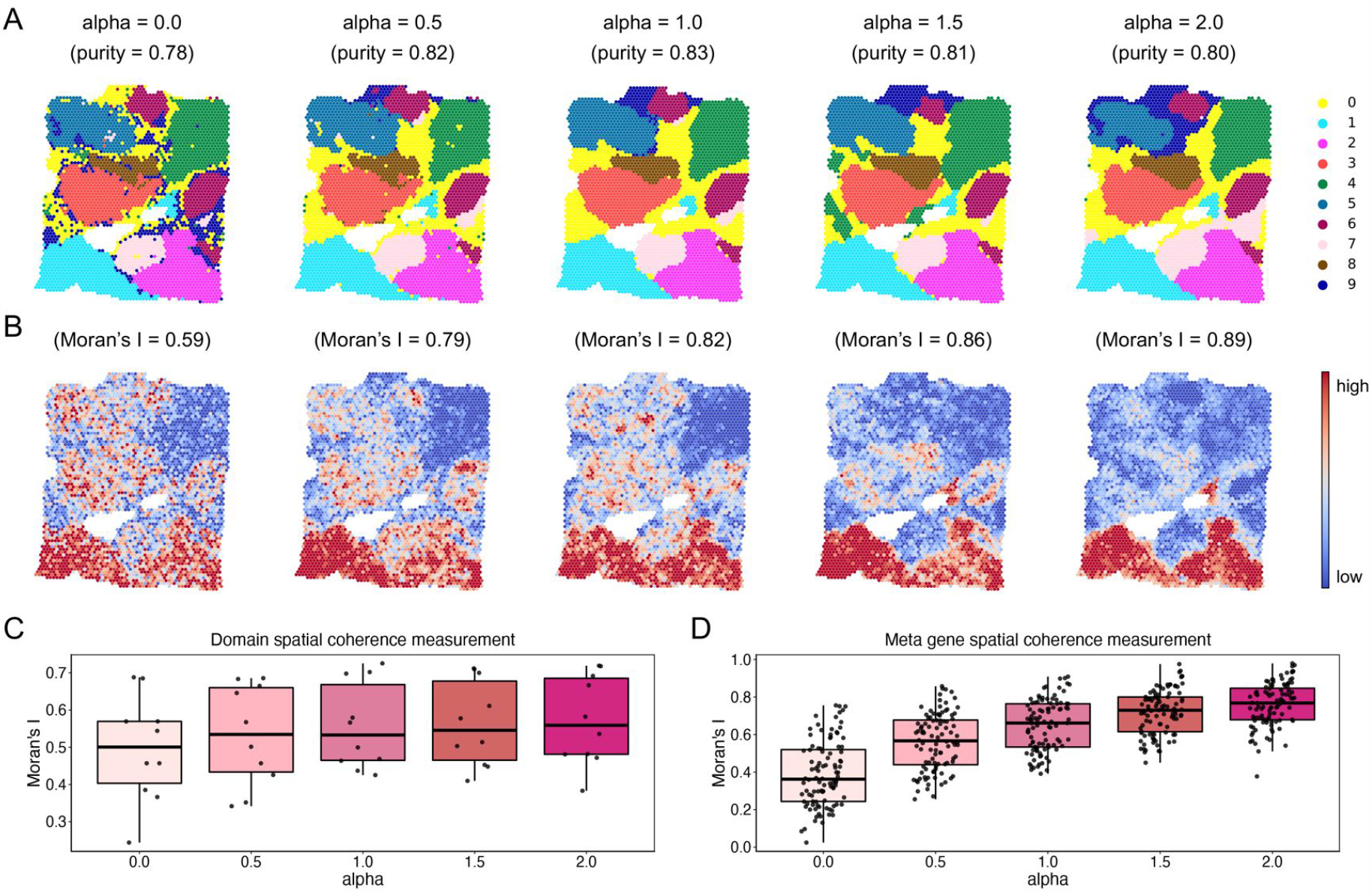
The illustrative analysis of flexible usage of spatial information on spatial domains and meta genes achieved by SpaSRL using the BC slice. (A) Spatial domains generated by SpaSRL under a variety of alpha settings. The identified spatial domains are distinguished using different colors and are shown on the spatial coordinates. The cluster purity is used to compare the similarity between the identified spatial domains and the ground truth annotations. (B) The spatial distribution of representative functional meta genes (focused on the bottom two spatial domains) identified by SpaSRL under a variety of alpha settings. Moran’s I measures the spatial autocorrelation of these functional meta genes. (C) Spatial coherence measurements of the identified domains from (A). (D) Spatial coherence measurements of the top 50 functional meta genes using Moran’s I statistics.

**Figure 4.**
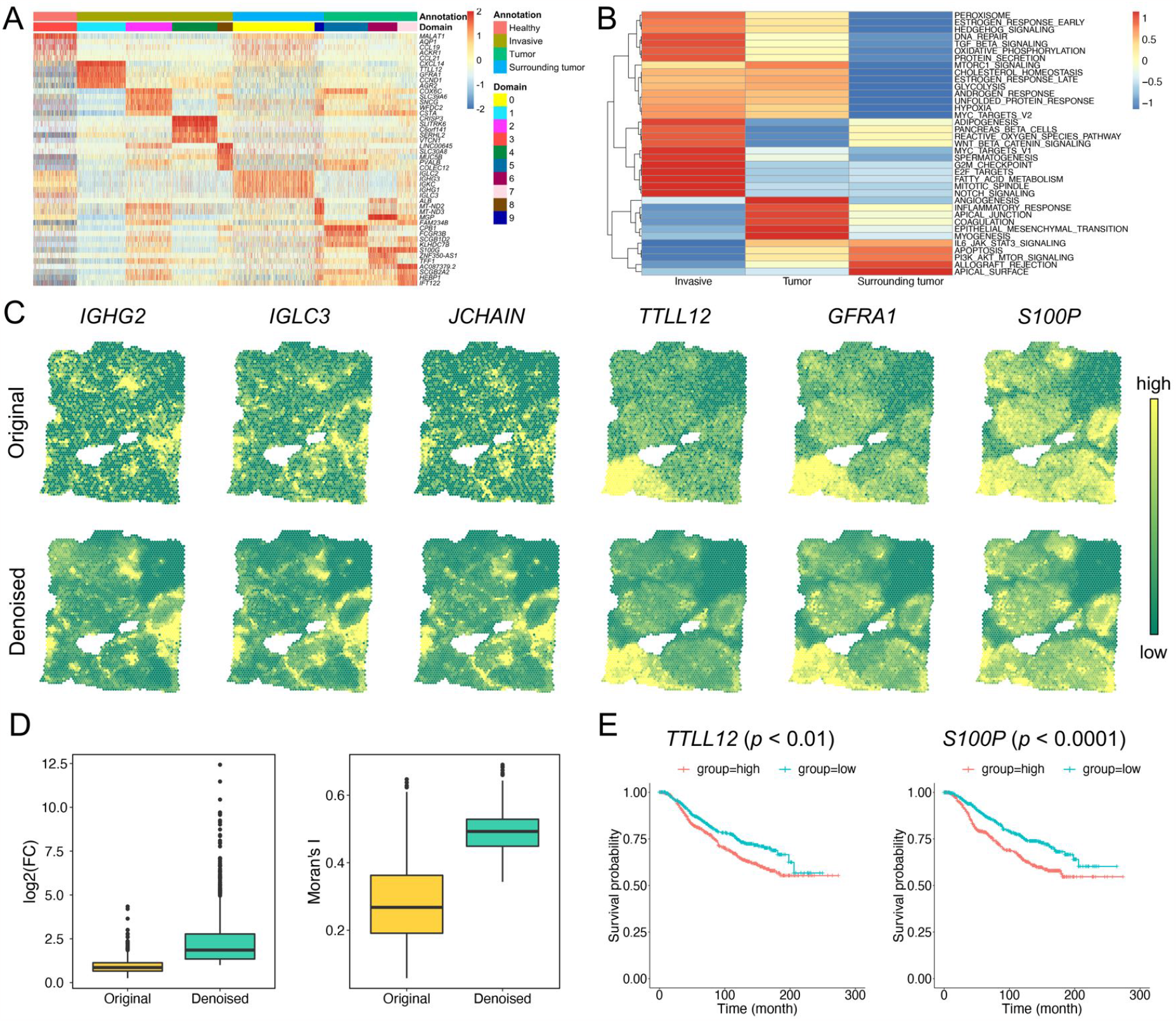
SpaSRL provides more biological insights into intratumor heterogeneity in BC sample. (A) The heatmap of original expression profiles of top 5 functional genes for each spatial domain identified by SpaSRL in Figure 2C. Each row of the heatmap indicates a gene and each column indicates a spot. The spots are labeled by histopathological annotations and SpaSRL assignments using column side colors. (B) The cancer hallmark enrichment of 357 functional genes in the Invasive, Tumor and Surrounding tumor regions. (C) Spatial expression of selected domain marker genes before (above) and after (below) data denoising. (D) The change of gene differential expression and spatial autocorrelation patterns before and after data denoising. FC: fold change of gene expression. (E) Survival analysis of the originally identified marker gene (i.e., *TTLL2*) and the newly identified marker gene (i.e., *S100P*).

### SpaSRL introduces a tunable strategy to achieve the flexible usage of spatial information

Next, we further clarified the competitive advantage of SpaSRL on integrating expression measurements and spatial information to improve the identification of spatial domains and meta genes with coherent expression and biological interpretation. Most current methods for SRT data directly constrain the models with spot spatial information, which facilitate the identification of expression patterns with spatial coherence but are more likely to overwhelm expression difference (Figure 2C and Supplementary Figure 3). To address this issue, SpaSRL provides a novel tunable spatially aware strategy to take account of transcriptional and spatial similarity by enhancing and decoding the shared information across spots (see Methods). In enhancing process, the tunable parameter alpha (*α*), which controls the shared expression between each spot and its surrounding neighbors, is used to adjust expression values in each spot (Figure 1A). Intuitively, when alpha is larger, the more shared information from spatial neighborhoods were aggregated in the enhancing process; while, during decoding process, spatially local similarity can occupy more in characterizing tissue structure and extracting spatial meta genes. Through such stepwise schema, SpaSRL transfers spatial correlation constraint between spots and genes, together with the flexible setting of alpha value, enabling the detection of spatial domains and functional (meta) genes with both spatial coherence and expression variability.

Here, we evaluated the effectiveness of our spatially aware strategy and validated the applicable range of alpha using BC slice. We varied alpha from 0 to 2 with increments of 0.5 to generate a series of enhanced profiles for evaluating the performance of identifying the functional meta genes and the spatial clustering (Figure 3). We computed the (1) cluster purity for evaluating accuracy of spatial domains (Figure 3A); and (2) Moran’s I statistics and Local Inverse Simpson’s Index (LISI) for measuring spatial coherence of spatial domains and functional meta genes (Figures 3C, 3D and Supplementary Figure 10). Based on these metrics, we found that SpaSRL identified spatial domains and functional meta genes with increasing spatial coherence as the alpha value became larger (Figures 3B-D and Supplementary Figure 10). However, the clustering purity exhibited a trend from rising to decline (alpha = 1.0 with the highest purity = 0.82, Figure 3A), indicating the varying consistency with histopathological annotation where the subtle biological differences might be missed if excessive spatial smoothing was implemented. The similar results can also be seen with the high-resolution Slide-seq V2 dataset (Supplementary Figure 11). Thus, these results show that how to effectively use spatial information in SRT model is critical to the rationale of clustering outcomes and gene-expression spatial distributions. By flexibly setting the alpha value, SpaSRL has great potential to reveal the biologically meaningful spatial regions and functional meta genes, adapting to more SRT technologies.

### SpaSRL provides more biological insights into intratumor heterogeneity on breast cancer

We had demonstrated SpaSRL can effectively dissect intratumor heterogeneity in BC as complementary to histopathological annotation (Figure 2C). In fact, SpaSRL can identify domain-specific functional genes (see Methods) and enhance the spatial expression patterns of individual genes, thus providing more insights to explore the molecular mechanisms underlying tumor heterogeneity.

In total, we obtained 357 spatial functional genes for all the identified domains (see methods) (Supplementary Figure 12). These spatial functional genes show high transcriptional specificity across spatial clusters and reveal distinct tumor microenvironments in the annotated cancer regions (Figure 4A). Then, performing cancer hallmark enrichment analysis (see Methods), we found that these spatial functional genes involved in different cancer-related biological processes, suggesting the potential cancer progression (i.e., surrounding tumor → tumor → invasive) in BC slice from the overall hallmark activation perspective (Figure 4B). These results indicated that SpaSRL could discover spatial functional genes with biological correspondence to heterogeneous tumor states.

Additionally, we validated the effectiveness of SpaSRL on denoising expression profiles to enhance or recover gene spatial expression patterns. After SpaSRL denoising, some spatial functional gene expressions (e.g., *IGHG2, IGHC3, JCHAIN, TTLL12, GFRA1* and *S100P*) appeared more spatially smoothed and with greater domain specificity on spots *in situ* (Figure 4C). The overall comparison of gene log fold change (LFC) and Moran’s I values quantifies the significant improvement of spatial expression coherence and biological specificity across domains brought by SpaSRL denoising (Wilcoxon signed-rank test *P* < 10^−16^ for LFC and *P* < 10^−13^ for Moran’s I, Figure 4D). We found 261 novel spatial functional genes in addition to the 96 genes that were ever identified in the original data under the same criteria (see Methods), indicating SpaSRL of potential to reveal new biological discoveries of disease. To further investigate this issue, we used two spatial domains (i.e., domain 0 in surrounding tumor region and domain 1 in invasive tumor region, Figure 2C) to display the LFCs of these spatial functional genes before and after denoising (Supplementary Figure 13), and observed the obvious enhancement of gene spatial expression for individual domains. Among the spatial functional genes for domain 1, 20 (out of 44) were validated to be the potential prognostic risk factors for breast cancer (Supplementary Table 1). These prognostic-related genes contained the originally and newly discovered genes. For example, *TTLL12* is an originally identified gene and ever reported to be positively correlated with poor prognosis in breast cancer [33] (Figure 4E). *S100P* is a new-found gene and proved as involved in the aggressive properties of breast cancer cells [34], which is upregulated in breast cancer and associated with poor prognosis (Figure 4E). These findings indicate that SpaSRL can distinguish intratumor heterogeneous regions but also can provide the comprehensive biological insights into the underlying heterogeneity by combing data denoising and spatial functional genes identification.

### SpaSRL identifies fine-grained mouse brain structures in 10x Visium datasets

We then applied SpaSRL to 10x Visium mouse brain sagittal sections (i.e., anterior and posterior samples) for comprehensive characterization of the fine-structured tissue architecture and region-specific functional genes. We took the hematoxylin and eosin (H&E) images of each dataset and the corresponding anatomical diagrams obtained from Allen Brain Atlas (ABA) as reference, and compared the anatomical regions with computationally generated domains by SpaSRL and other competing methods.

For the anterior slice, SpaSRL distinguished the domains largely consistent with the ABA and H&E references, including the layered cortical structures of five cerebral cortex (CTX) domains (i.e., domain 1, 2, 5, 6 and 7) and fiber tract (i.e., domain 3), and a subtle region of the lateral ventricle (VL) section (i.e., domain 13). While other benchmarking methods failed to localize the fine structures or identified fewer sections (Figure 5A and Supplementary Figure 14). Additionally, BayesSpace can also identify the layered domains (i.e., CTX and fiber tract), but SpaSRL’s separation showed better transcriptional specificity on the layer known marker genes (from outer to inner layers: *Ptgds, Rasgrf2, Stx1a, Myl4, Nptx1* and *Plp1*) (Figure 5B and Supplementary Figure 15). For the VL section, SpaSRL also detects its specific functional meta gene (Figure 5C), where the top weighed genes are *Enpp2* and *Ttr* (Figure 5D), two marker genes of choroid plexus epithelial cell type which is enriched in VL region [35]. Thus, SpaSRL can detect the fine-grained brain structures and identify the biologically informative genes that underlie the corresponding spatial domains.

**Figure 5.**
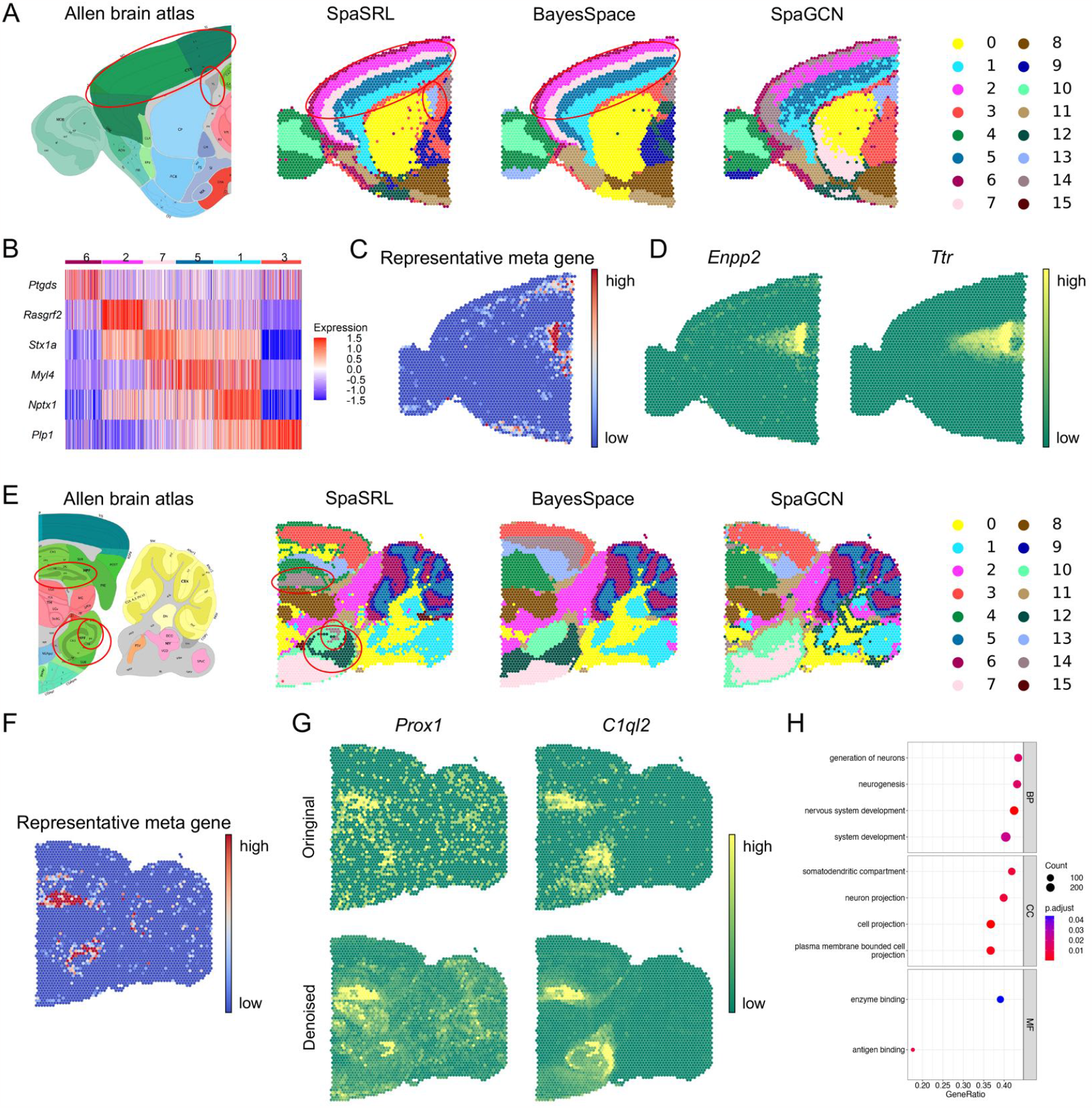
SpaSRL identifies tissue structures and functional genes/meta genes in mouse brain sagittal anterior (*n* = 3,696 spots) (A-D) and posterior (*n* = 3,353 spots) (E-H) slices. The corresponding anatomical definitions obtained from the Allen Mouse Brain Atlas (First image in A and E) are shown as references. The identified spatial domains by all the involved approaches are illustrated on the spatial coordinates and distinguished using different colors without anatomical correspondence. Fine anatomical regions, for example CTX, fiber tract, VL sections in (A) and DG, CA sections in (E) are marked by red circles on reference images and computational results (if any exists). (B) The original expression heatmap of known marker genes separates the CTX and fiber tract layers identified by SpaSRL. SpaSRL CTX layers (from outer to inner) contain domains 6, 2, 7, 5 and 1; fiber tract is the domain 3. (C) The representative functional meta gene identified by SpaSRL to characterize VL section. (D) The original spatial expression of the top two weighted functional genes in the representative functional meta gene from (C). (F) The representative meta gene identified by SpaSRL to characterize two separated DG sections. (G) Spatial expression of the two marker genes of DG sections before and after denoising. (H) The GO enrichment of the representative meta genes from (C). The size indicates the number of functional genes enriched in each GO term and the color indicates the statistical significance (adjusted by false discovery rate [FDR]) of enrichment in each GO term. The terms are grouped by GO subontology. BP, Biological Process; CC, Cellular Component; MF, Molecular Function.

For the posterior slice, only SpaSRL recognized the dentate gyrus (DG) (i.e., domain 14), and cornu ammonis (CA) (i.e., domain 12) sections of the hippocampus region, just as the H&E stained shape on original image and ABA reference (Figure 5E and Supplementary Figure 16). On the slice, the domain 14 contains two separated regions that can be clearly highlighted by the SpaSRL obtained meta gene (Figure 5F) and two DG marker genes (i.e., *Prox1* [36] and *C1ql2* [37]). Moreover, SpaSRL greatly improved the marker gene spatial expression and specificity on the denoised profiles (Figure 5G). We then extracted the spatial functional genes of this domain and performed functional enrichment analysis (see Methods) (Supplementary Figure 17). We found these spatial functional genes were enriched in many biological functions related to hippocampus DG region, e.g., neurogenesis, generation of neurons and nervous system development [38] (Figure 5H). These results indicate that SpaSRL not only can detect subtle spatial biological signals but also can effectively decode spatial expression patterns from spot and gene perspectives with correspondent associations and biological interpretations. The effectiveness of SpaSRL at identifying fine-grained tissue structures and enhancing the spatial expression of functional (meta) genes enables the potential applications on high-resolution SRT platforms with high dropout rates.

### SpaSRL reveals spatial expression landscape in Slide-seqV2 cerebellum dataset

We next verified that SpaSRL can obtain spatial domains at finer-grained level or even distinguishing individual cell types based on a mouse cerebellum Slide-seqV2 dataset. Here we leveraged the anatomical diagrams and cell-type related marker gene sets (from Allen brain atlas and Cable et al.’s previous work [39]) for cluster annotation and comparison analysis. SpaSRL identified 16 clusters in total, which not only accurately correspond to the cerebellum anatomical structures or cell types, but also have better spatial coherence than clusters obtained by SpaGCN (other methods not involved due to model limitations) (Figure 6A, B and Supplementary Figures 18-20).

**Figure 6.**
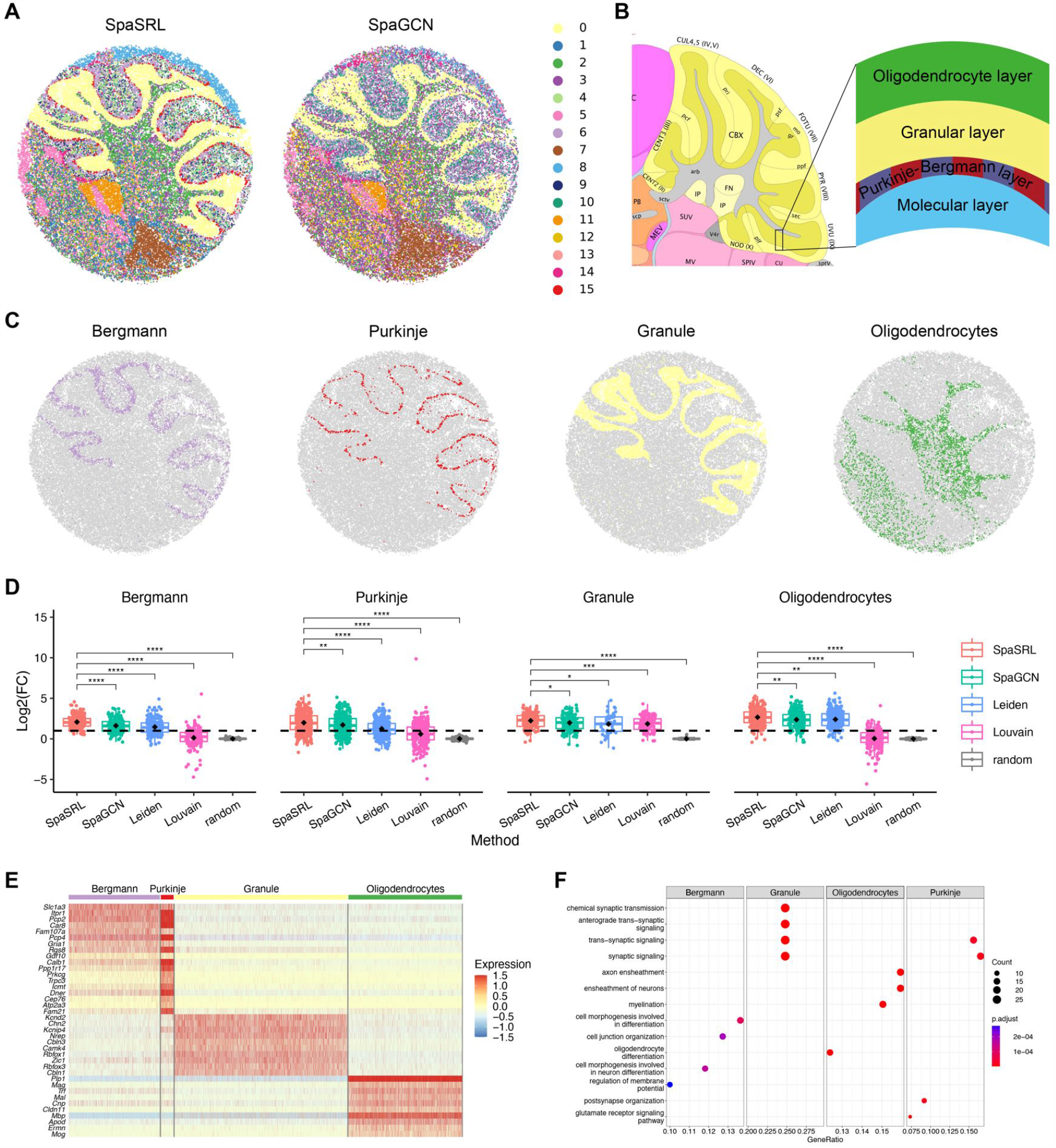
SpaSRL reveals spatial expression landscape in Slide-seq cerebellum data (n = 39,496). (A) Each bead on the spatial coordinates is colored by the cluster assignments of SpaSRL and SpaGCN. (B) The regional structure diagrams from Allen brain atlas and Cable et al.’s previous work [39] are used to directly compare and illustrate clustering results. (C) Individual loadings of four layers of cerebellum identified by SpaSRL. (D) Boxplots of expression fold change (FC) of marker gene sets for the selected four cluster assignments identified by SpaSRL and other methods. The black dot in boxes represents average log2(FC) of all involved markers. Dashed line indicates the log2(FC) value of 1. Difference of log2(FC) distributions between SpaSRL and other methods are quantified by Wilcoxon signed-rank test, and the significance is shown if the FDR-adjusted *P*-value is lower than 0.05. **: *p*-value < 0.01, ***: *p*-value < 0.001, ***: *p*-value < 0.0001. (E) The original expression heatmap of the top 10 functional genes for the four clusters in (C) identified by SpaSRL. (F) The bubble plot of GO BP enrichment results of all the functional genes for each cluster in (C). The dot size indicates the number of functional genes enriched in each GO term and the color indicates the FDR-adjusted significance of enrichment in each GO term.

More notably, SpaSRL succeeded to figure out the finer-grained layered organization (from the outer layer of Bergmann cells to the inner of oligodendrocytes) of cerebellum (Figures 6A-C). Based on the locally enhanced expression *X*, we calculated the LFCs of the marker genes from RCTD [32] to quantify the clustering performance of each method and high average LFC value obviously indicates high biological specificity between the identified clusters. As expected, these computationally obtained clusters appeared to be more biologically meaningful than random separations (Figure 6D, Wilcoxon signed-rank test, *P* < 10^−30^). While there are still significant differences between the involved methods. SpaSRL performed substantially better to distinguish these structures or cell types (Figure 6D), especially to separate the colocalized Purkinje and Bergmann cells [39] with significantly higher transcriptional specificity (Wilcoxon signed-rank test, *P* < 10^−3^). As for Louvain, the clusters (granule excluded) were less biologically coherent (Wilcoxon signed-rank test, *P* < 10^−20^) with high FC scores in only a few markers. These results suggest that SpaSRL could better unveil the organizational landscape of complex tissues in higher resolution spatial dataset.

Additionally, we used the spatial functional genes of these layered structures to further explore their biological differences (see methods) (Supplementary Figure 21). The heatmap of the top spatial functional genes illustrates the transcriptional similarities between these cell types (Figure 6E) where Bergmann and Purkinje cells share some marker genes due to their spatial colocalization (Figure 6A). While, we further made functional enrichment analysis, and found the identified spatial functional genes were involved in different biological processes which can clearly distinguish all these structures (Figure 6F). For example, Purkinje and granule cells are involved in sending nerve signals in cerebellar cortex [40], while some synaptic signaling related biological functions (i.e., synaptic signaling and trans-synaptic signaling) are enriched for both cell types. Oligodendrocytes are the myelinating cells of the central nervous system [41], and the processes of myelination and ensheathment of neurons are enriched using the identified oligodendrocytes related genes. In summary, for higher resolution spatial data, SpaSRL can reveal spatial expression landscape and can effectively detect functional gene sets to explain the biological spatial heterogeneity of tissue architecture.

## Discussion

Spatial mRNA measurements provide new perspectives to define biological heterogeneities of tissues and diseases under spatial context. The fundamental issue of deciphering tissue heterogeneity is to accurately capture the relationship between spots and the functional genes with spatial coherence and co-expression. However, the current methods still focus on the individual task and there is a great challenge to join the two tasks into a unified model while flexibly incorporating spatial information. In this work, SpaSRL, was designed as such a scalable framework that accounts for these requirements. SpaSRL provides a computationally efficient tool to obtain the spot-spot relations and the functional genes with flexibly spatial correlation constraint, which are then used for accurate downstream analysis, including spatial domain detection, spatial functional genes/meta genes identification and data denoising. The superiority of SpaSRL not only is reflected on the accurate identification of spatial domains and functional genes on multiple datasets of various technologies, i.e., 10x Visium, Slide-seqV2, but also can control the usage of spatial information to impact the identification of spatial domain and functional genes. In particular, the application on breast slices and brain slices demonstrated that SpaSRL reveals the tissue structures and the corresponding spatial functional genes for interpreting tissue heterogeneity, suggesting that SpaSRL is more suitable for deciphering complex spatial expression landscape in SRT study.

The effective usage of spatial information is key to the superiority of SpaSRL in SRT study. Specifically, SpaSRL first uses spatial information to enhance the shared information between neighboring spots. Then, SpaSRL builds a novel self-representation model to decode the shared expression between spots from the enhanced data of the low-dimensional space and the original space. The stepwise approaches of SpaSRL take spatial information as a soft constraint to impact the spatial coherence of the spatial domain and functional meta genes, and to mitigate SRT data noise or bias. Additionally, the novel self-representation learning model achieves dimension reduction and spatial domain detection into a unified model, which makes that the spatially aware strategy of SpaSRL transfers spatial correlation constraint between two tasks. Thus, compared to the current methods, SpaSRL not only flexibly controls the usage of spatial information but also improves the identification of biologically informative spatial domain and functional genes by leveraging spatial information.

Although SpaSRL provides the joint analysis of the spatial domain detection and the functional genes identification, SpaSRL still performs the individual gene analysis and cannot track which functional gene network module contributes to each spatial domain, which is a limitation of SpaSRL for application to the in-depth analysis of biological interpretations. Certainly, most methods are compromised to use a separate manner: first to detect spatial domain and subsequently to infer gene networks (or vice versa). However, the splitting solution may cause untraceable biological variabilities, even leading to biased outcomes. Therefore, further study of the joint analysis of domains and gene networks under spatial context is warranted.

## Data availability

The human dorsolateral prefrontal cortex (DLPFC) datasets are available in the spatialLIBD package (http://spatial.libd.org/spatialLIBD). The Mouse Brain Sagittal, Invasive Ductal Carcinoma and Human Breast Cancer datasets are available at 10x Genomics website (https://www.10xgenomics.com/resources/datasets). The Slide-seqV2 data is available at https://singlecell.broadinstitute.org/single_cell/study/SCP815.

## Code availability

Python source code of SpaSRL, under the open-source BSD 3-Clause license, is available at https://github.com/zccqq/SpaSRL. The documentation website provides the installation guide, tutorials, and API references, which is available at https://spasrl.readthedocs.io/. SpaSRL is also published as a Python package named ‘spasrl’ on Python Package Index (PyPI) at https://pypi.org/project/spasrl/ and can be directly installed via the pip installer.

## ACKNOWLEDGEMENT

We thank the anonymous reviewers for useful suggestions. Q.S. and C.Z conceived and designed the framework and the experiments. L.W. and L.X. performed the experiments. L.W. and H.D. developed the Python package and documentation website of the framework. L.W., Q.S. and C.Z analyzed the data and wrote the paper. Q.S. and C.Z revised the manuscript.

## KEY POINTS

1. SpaSRL is a spatially aware self-representation learning method that unify the identification of spatial domains and functional (meta-) genes, relying on a joint model to optimally transfer spatial correlation constraint between the two tasks.
2. SpaSRL provides a novel tunable strategy to achieve the flexible integration of spatial information via an enhancing-decoding schema as complementary to the current usage of spatial information.
3. SpaSRL is a user-friendly and computationally efficient Python tool that can be scalable for diverse SRT platforms.

### Chuanchao Zhang

received the PHD from Wuhan University, Wuhan, China, in 2017. He is currently an assistant research fellow in Key Laboratory of Systems Health Science of Zhejiang Province,

Hangzhou Institute for Advanced Study, University of Chinese Academy of Sciences, Chinese Academy of Sciences, Hangzhou 310024, China. His current research interests include machine learning, deep learning, single-cell transcriptomics and spatial transcriptomics.

### Xinxing Li

is a master student in College of Informatics, Huazhong Agricultural University, Wuhan, China. Her current research interests include single-cell transcriptomics and spatial transcriptomics.

### Wendong Huang

is a master student in College of Informatics, Huazhong Agricultural University, Wuhan, China. His current research interests include single-cell transcriptomics and spatial transcriptomics.

### Lequn Wang

is pursuing his PhD degree in the Center for Excellence in Molecular Cell Science. His current research interests include bioinformatics, machine learning, single-cell multi-omics and spatial transcriptomics.

### Qianqian Shi

received the PHD from Shanghai Institute of Biological Sciences, University of Chinese Academy of Sciences, Chinese Academy of Sciences, China, in 2017. She is currently an associate professor at College of Informatics, Huazhong Agricultural University, Wuhan, China. Her current research interests include machine learning, deep learning, network biology, computational biology, single-cell transcriptomics and spatial transcriptomics.

